# PinAPL-Py: A comprehensive web-application for the analysis of CRISPR/Cas9 screens

**DOI:** 10.1101/147462

**Authors:** Philipp N. Spahn, Tyler Bath, Ryan J. Weiss, Jihoon Kim, Jeffrey D. Esko, Nathan E. Lewis, Olivier Harismendy

## Abstract

**Background:** Large-scale genetic screens using CRISPR/Cas9 technology have emerged as a major tool for functional genomics. With its increased popularity, experimental biologists frequently acquire large sequencing datasets for which they often do not have an easy analysis option. While a few bioinformatic tools have been developed for this purpose, their utility is still hindered either due to limited functionality or the requirement of bioinformatic expertise.

**Results:** To make sequencing data analysis of CRISPR/Cas9 screens more accessible to a wide range of scientists, we developed a Platform-independent Analysis of Pooled Screens using Python (PinAPL-Py), which is operated as an intuitive web-service. PinAPL-Py implements state-of-the-art tools and statistical models, assembled in a comprehensive workflow covering sequence quality control, automated sgRNA sequence extraction, alignment, sgRNA enrichment/depletion analysis and gene ranking. The workflow is set up to use a variety of popular sgRNA libraries as well as custom libraries that can be easily uploaded. Various analysis options are offered, suitable to analyze a large variety of CRISPR/Cas9 screening experiments. Analysis output includes ranked lists of sgRNAs and genes, and publication-ready plots.

**Conclusions:** PinAPL-Py helps to advance genome-wide screening efforts by combining comprehensive functionality with user-friendly implementation. PinAPL-Py is freely accessible at http://pinapl-py.ucsd.edu with instructions, documentation and test datasets. The source code is available at https://github.com/LewisLabUCSD/PinAPL-Py

## Background

Genetic screens using pooled CRISPR/Cas9 libraries are functional genomics tools that are becoming increasingly popular throughout the life sciences. Using these screens, researchers are able to find novel molecular mechanisms, better understand complex cellular systems, or find new drug targets [1–3]. Analysis of the raw sequencing output of these screens is a non-trivial task, given the size and diversity of these datasets. MAGeCK [4] was the first bioinformatic workflow specifically aimed at analysis of CRISPR/Cas9 screens and has been used in numerous genome-wide screening studies since. However, while being a standard solution in the field, MAGeCK has to be run from a command line, and requires manual definition of the position of the 20 bp single guide RNA (sgRNA) sequence for proper read identification. It, thus, requires familiarity with working from a command-line, basic handling of raw fastq files, as well as knowledge of the read sequence composition, all of which might constitute obstacles to many biologists. BAGEL [5] is an alternative command-line workflow, powerful in analyzing gene essentiality screens, but inapplicable to other screening experiments, such as treatment resistance screens or reporter-based screens. caRpools [6] is a R-package offering multiple analysis options, but installation and execution require proficiency with the R platform, and support of the package seems to have been discontinued. Finally, the recently developed CRISPRcloud [7] is an analysis workflow that runs as a web-service, thus offering superior ease of use. However, CRISPRcloud requires manual definition of a fixed 20 bp window to extract the sgRNA sequence from each read file making the workflow incompatible with cases where the position of the sgRNA sequence varies between reads sequenced on the same instrument, a protocol referred to as read staggering. Read staggering is a common technique and recommend by many sequencing facilities to increase sequencing yield as it mitigates the low base diversity problems in the initial sequencing cycles of PCR amplicons libraries like the ones generated in CRISPR/Cas9 screens [8–10] (Fig. 4A). As a consequence, sequencing analysis of CRISPR/Cas9 screens still remains challenging for most laboratories since prior solutions are either hard to access for non-bioinformatic experts, or might not provide sufficient support for the datasets generated. The web-service we present, PinAPL-Py, addresses these issues by providing a comprehensive analysis workflow running through an intuitive web-interface that supports a broad class of CRISPR/Cas9 screening experiments as well as staggered reads. This facilitates standardized, reproducible data analysis that can be carried out directly by the scientists conducting the experiments.

## Implementation

### Workflow description

PinAPL-Py is designed to analyze a generic layout of a CRISPR/Cas9 screening experiments where a control group is compared to one or more conditions which can, for example, refer to cells exposed to a chemical compound, samples taken at different time points, or cells expressing a fluorescent reporter and sorted by FACS (Fig 1A). PinAPL-Py is written in Python 2.7 and implements well-established methods to provide a fully automated workflow, taking the user from the raw sequence files to a list of candidate sgRNAs and genes while requiring only minimal manual input. In particular, the workflow comprises the following steps (Fig. 1B):

**Fig 1:**
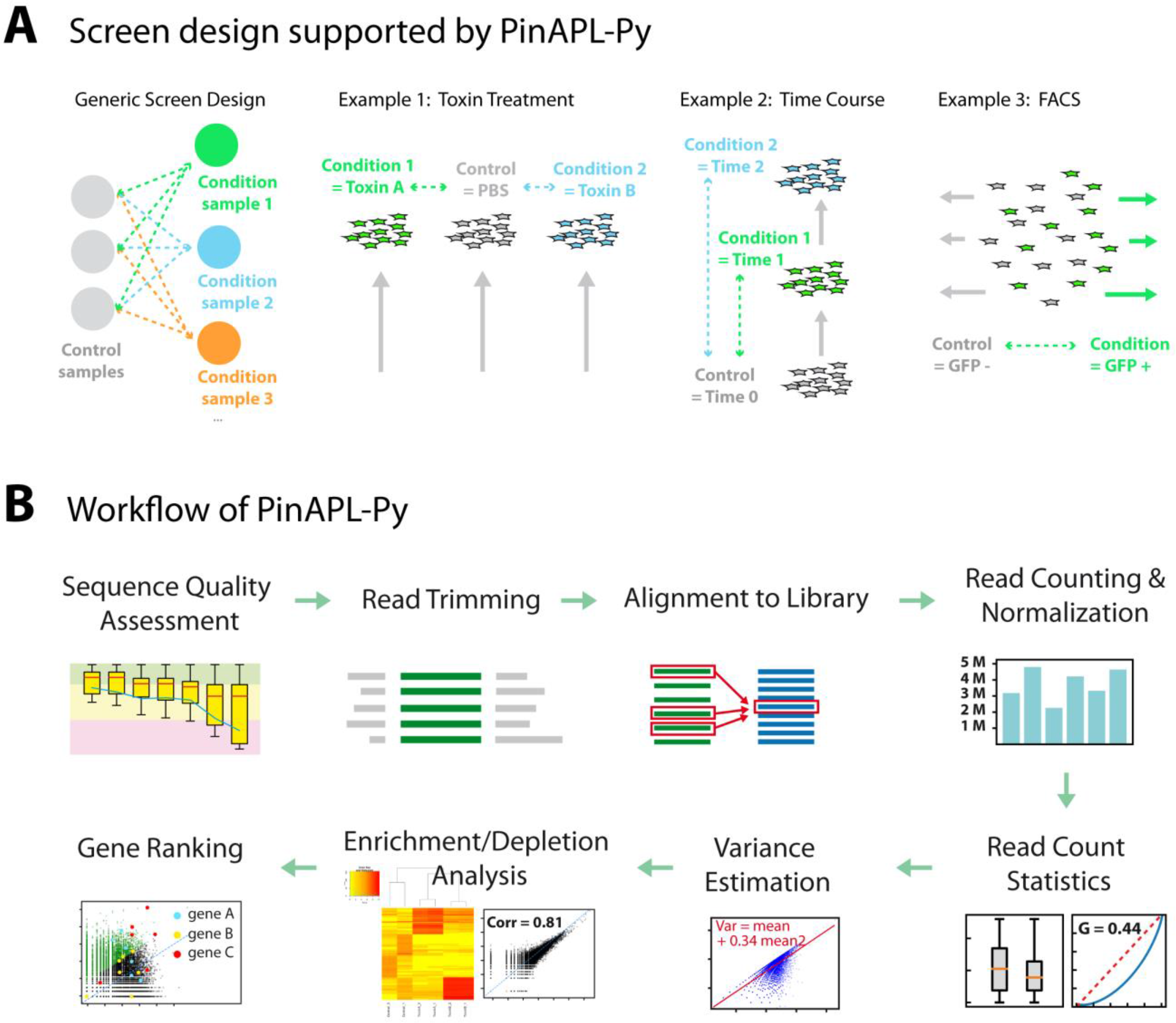
PinAPL-Py’s scope and methods. **(A):** Examples of screen designs, compatible with PinAPL-Py. PinAPL-Py is designed to analyze a generic screen in which a single control (possibly represented by several replicates) is to be compared to a number of “conditions” which can take various forms, depending on the context of the experiment. 1-3 show common examples, but are not an exhaustive list. Comparisons can either look for sgRNAs/genes being depleted or enriched relative to the control. **(B):** Overview of the analysis workflow implemented in PinAPL-Py. For details see text.

#### Sequence quality check

Sequence quality assessment is carried out using fastqc (http://www.bioinformatics.babraham.ac.uk/projects/fastqc/), which analyzes sequence composition, sequencing quality, and read depth.

#### Read trimming

Before alignment of the sequencing reads to the library, the reads are processed with the *cutadapt* tool to remove known adapter sequences located 5’ of the 20 bp sgRNA sequence [11]. These adapters are library-specific sequences, generated in the sgRNA cloning process, and are often unknown to the user. Adapter information for the 19 most commonly used libraries is stored in PinAPL-Py so that the user only needs to manually provide their adapter sequence if working with a custom library. By default, *cutadapt* is run with an alignment error tolerance of 0.1, and minimal required sequence length of 20 bp, after trimming. After removing the 5’ adapter, *cutadapt* trims the remaining read to exactly 20 bp to retain only the sgRNA sequence.

#### Read alignment

Read alignment is carried out using *Bowtie2* [12] in the –local mode. *Bowtie’s* alignment parameters (seed length (-L), seed number (-N), and seed interval function (-i)) are set to values producing optimal results for 20 bp sgRNA sequences, but can be changed on the configuration page. A technical description of these parameters is available in the *Bowtie2* manual (http://computing.bio.cam.ac.uk/local/doc/bowtie2.html#what-is-bowtie-2). After completion, alignments are processed based on the best (=primary) and second-best (=secondary) alignment scores achieved for each read (referred to ‘AS’ or ‘XS’, respectively, in the *Bowtie2* SAM output), and categorized as either “unique”, “tolerated”, “ambiguous”, or “failed”. An alignment is considered failed if either no primary score is reported by *Bowtie2*, or if the primary score reported is below the critical matching threshold (set to perfect matching in the default setting, but adjustable on the configuration page). An alignment is considered ambiguous if the difference between primary and secondary alignment is below a critical ambiguity threshold (adjustable on the configuration page). If this difference is above the threshold, the read is mapped to the primary match and classified as “tolerated”. If a secondary score is not reported (and the primary score satisfies the matching threshold), the alignment is considered unique. Only unique and tolerated reads are kept for further processing, while failed and ambiguous reads are discarded.

#### Read counting and normalization

The abundances of retained reads are quantified for individual sgRNAs, and also for each gene by summing all reads belonging to all of the sgRNAs targeting that gene. To remove noise, the user can define a critical count cutoff (adjustable on the configuration page), below which read counts (given in counts per million reads) are set to 0. Read counts are normalized by a method of choice, specified on the configuration page. Available options are counts per million reads (cpm), “total” normalization (read counts, divided by the number of total read counts in the sample and multiplied by the mean total read count across all samples of the experiment), or using the “size-factor” method which uses a median ratio to correct for different total read counts [4,13].

#### Analysis of read count distributions

For each sample, descriptive statistics of normalized read counts are computed and reported on the results page as boxplots, histograms and various measures, such as median, standard deviation, maximum, minimum, sgRNA/gene representation, and the Gini coefficient.

#### Read count variance estimation

In order to generate a candidate list of individual sgRNAs showing either the most significant enrichment or depletion (compared to the control samples), PinAPL-Py models normalized read counts as following a negative binomial distribution [14], similarly to other read count analysis workflows [4,15,16]. For this, the average (normalized) read count μi and sample variance 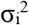 for each sgRNA *i* is computed across all control replicates. The dispersion parameter *D* of the negative binomial distribution is then estimated via linear regression since σ^2^ = μ + D μ^2^ holds in a negative binomial model. With this estimated dispersion, a model variance 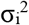 is computed for each sgRNA. Thus the read count of each sgRNA *i*, under the control condition, is modeled to follow a negative binomial distribution with parameters NB(μi, 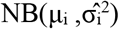).

#### sgRNA enrichment / depletion analysis

sgRNA counts from each treatment sample are then evaluated on the basis of the read count distribution, estimated in the previous step. In particular, for each normalized read count ci from a treatment sample, the fold-change to the control mean is taken, and a p-value is computed from NB(μi, 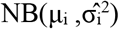) to test either for ci > μi (in an enrichment screen) or ci < μi (in a depletion screen). Finally, these p-values are adjusted for multiple hypothesis tests using the Benjamini-Hochberg FDR procedure. Note that for the computation of p-values, PinAPL-Py requires the user to specify at least two control samples (only fold-change is reported otherwise). The results are returned to the user as sortable tables on the results page, and as spreadsheets after download. In addition, PinAPL-Py produces scatterplots of normalized counts for each treatment sample versus the control averages on which sgRNAs of individual genes can be interactively highlighted on the results page. Results of the sgRNA enrichment/depletion analysis are reported for each treatment sample separately, and correlation analysis is carried out for samples representing replicates of the same treatment.

#### Sample clustering

Unsupervised clustering analysis of all samples is done based on either the most variable or most abundant/depleted (depending on screen type) sgRNAs throughout the dataset. For most variable sgRNAs, the *n* sgRNAs with the highest variance across all samples are extracted (*n*=25 by default, adjustable on the configuration page). For highest abundance/depletion, the *n* sgRNAs with the highest/lowest read counts are extracted for each sample, and the union set of all these sgRNAs is used for the clustering. Clustering is done through the *gplots* package in R.

#### Gene ranking

From the sgRNA enrichment/depletion analysis described above, a gene ranking is derived by combining the fold-change data from all sgRNAs targeting a single gene through one of two possible methods:

- *Adjusted Robust Rank Aggregation (αRRA):* This option follows the αRRA procedure described previously [4]. Individual sgRNAs are required to show enrichment/depletion at significance level of *P_0_ = 5.10^−5^* (by default) in order to be taken into account for computation of the gene ranking score. Genes having no significant sgRNAs are given a score of 1.0; p-values for the gene ranking score are computed based on randomly assigning sgRNAs to target genes and running permutations (Np = 1,000 by default) to estimate the empirical distribution function.
- *STARS:* This option calls the STARS function to compute gene ranking scores and p-values based on a binomial model (http://portals.broadinstitute.org/gpp/public/software/index) Technical details are provided in the original publication [17].

### Benchmarking & Quality control

#### Sequencing data

Datasets for comparisons with MAGeCK/CRISPRcloud were taken from CRISPR/Cas9 screening experiments in our laboratory (unpublished). In these, an A375 cell population was transduced with the full human GeCKO library at ~100x coverage and split into treatment and control samples. Treatment samples were incubated with a toxin at ~90% lethal dose. Control samples were incubated with PBS. After 4 days, cells were harvested, sgRNA sequences were extracted with nested PCR [10], and sequenced at a minimum depth of 4 million reads per sample.

#### Sequencing data analysis

PinAPL-Py was run with gene ranking set to “αRRA” with 1,000 permutations. Matching threshold was set to 40 (perfect match) with ambiguity threshold set to 2 (Fig 2). Read count normalization was set to “total”. All other parameters were left at default. To generate the ranked list of top 25 genes (Fig. 3E), the gene list was sorted on two levels: First by αRRA FDR (low to high), and second by –log αRRA score (high to low).

**Fig 2:**
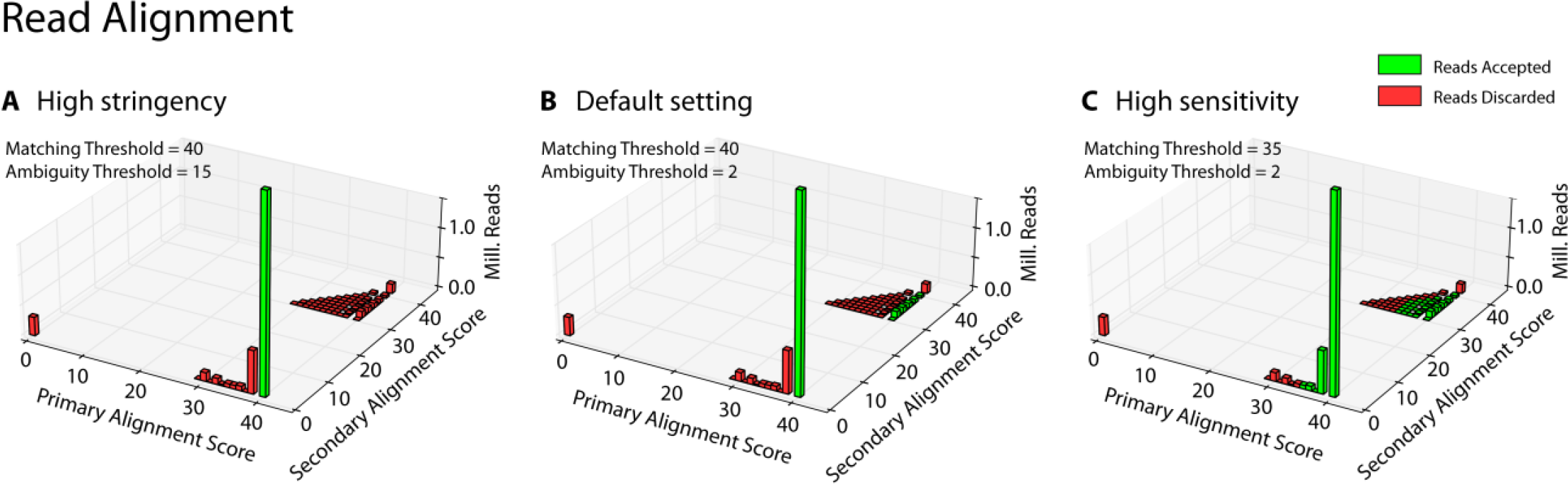
Read alignment optimization. Barplots of the primary (best matching library entry) and secondary alignment score (second-best matching library entry) achieved by each read. By adjusting the matching threshold, the user can include reads with non-perfect matches to increase sensitivity. By adjusting the ambiguity threshold, the user can control to what degree the algorithm should tolerate reads matching multiple library entries. Reads located on the diagonal match multiple library entries equally well (primary score = secondary score) and are discarded by PinAPL-Py’s default setting (As an example, the popular Human GeCKO library contains almost 4,000 sequences matching multiple target genes). **(A):** High stringency setting requires perfect matching and allows no (not even low) matching to another library sequence. **(B):** The default setting requires perfect matching, but also accepts reads if they have a second-best match in the library, as long as the second-best score is lower than the primary score. **(C):** High sensitivity setting accepts reads with a less than perfect sequence match, e.g. when accounting for less than perfect sequence quality.

**Fig 3:**
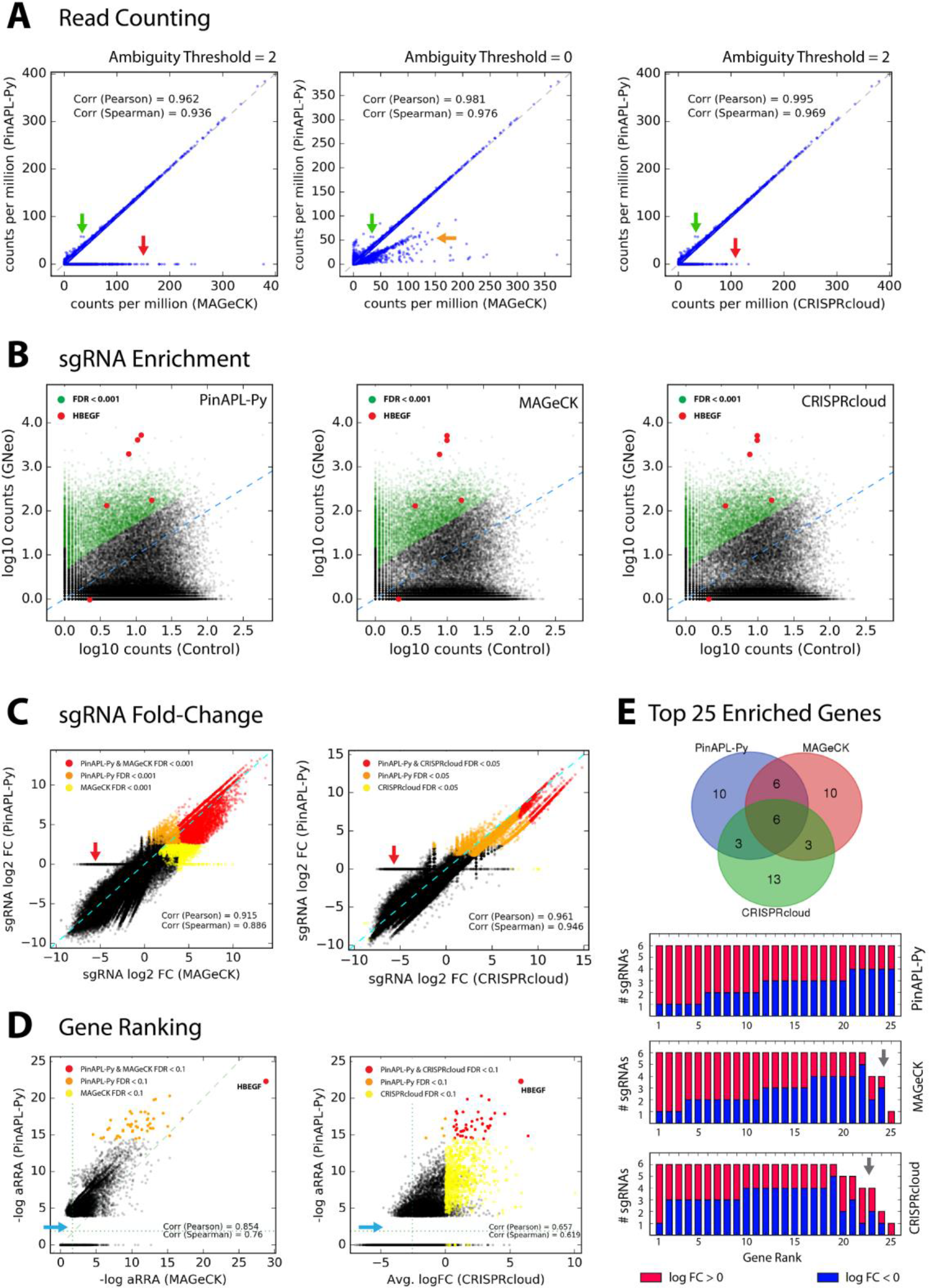
Comparison to other tools. **(A):** Read counts for each sgRNA obtained by PinAPL-Py (y axis) and MAGeCK (x-axis, left and middle panel) or CRISPRcloud (x-axis, right panel). The horizontal line of points at y=0 (red arrows) reflects PinAPL-Py’s rejection of reads that map equally well to multiple library entries (see Fig 2). When reducing the ambiguity threshold to 0, the horizontal line disappears as reads become randomly assigned to one of its matching library entries (middle panel, orange arrow). sgRNAs yielding higher counts in PinAPL-Py (green arrow) are due to failed alignments in MAGeCK/CRISPRcloud caused by indel frameshifts 5’ of the sgRNA sequence. **(B):** Scatterplots showing log10 (normalized) read counts for each sgRNA from the treatment sample (y-axis) versus control samples average (x-axis) (Left: PinAPL-Py, middle: MAGeCK, right: CRISPcloud). sgRNAs showing significant enrichment marked in green. sgRNAs targeting *HBEGF* shown in red. **(C):** log2 fold-changes (treatment relative to control average) for each sgRNA, compared between PinAPL-Py and MAGeCK (left) and PinAPL-Py and CRISPRcloud (right). sgRNAs yielding significant enrichment in both (red) or either (orange or yellow) method are indicated. The horizontal line of points at y=0 (red arrows) is a consequence of PinAPL-Py’s rejection of ambiguous reads (see A). **(D):** Gene ranking scores, compared between PinAPL-Py (αRRA) and MAGeCK (αRRA) (left), and PinAPL-Py (αRRA) and CRISPRcloud (average log2 fold-change). Genes yielding significant enrichment in both (red) or either (orange or yellow) method are indicated. The point gap on the low end of PinAPL-Py’s score range (blue arrows) is caused by its procedure to assign genes with no significant sgRNAs an αRRA score of 1.0 **(E):** Venn diagram of the top 25 genes, obtained by each method. Barplots below show the number of sgRNAs for each of the top 25 genes, marked red for enrichment (log2 fold-change > 0) and blue for depletion (log2 fold-change < 0). The grey arrow points at genes being represented by fewer than 6 sgRNAs in the library.

MAGeCK v0.5.6 was run following the instructions on sourceforge (https://sourceforge.net/p/mageck/wiki/Home/#usage). Normalization was set to “total”. To generate the ranked list of top 25 genes (Fig. 3E), the list was sorted on two levels: first by FDR (“pos|fdr”, low to high), and second by –log αRRA score (“pos|score”, high to low).

CRISPRcloud (http://crispr.nrihub.org/) was run in the “Survival and Drop-out” mode, with analysis algorithm set to “DESeq2” (since this option is most comparable to the negative binomial model implemented in PinAPL-Py and MAGeCK). To generate the ranked list of top 25 genes (Fig. 3E), the gene list was sorted on three levels: 1: FDR (low to high), 2: hit ratio (HR, high to low), and 3: average sgRNA log10 fold-change (AVGFC, high to low). Venn diagrams were generated at http://bioinformatics.psb.ugent.be/webtools/Venn/. For the comparisons between tools in Fig. 3, a dataset without staggers in the reads was used.

## Results & Discussion

### User input

To start a PinAPL-Py session, the user enters a project name on the home screen at http://pinapl-py.ucsd.edu. Optionally, an email address can be entered to avoid data loss in case of an unintentional browser shutdown and to receive notification upon completion of the analysis. Sequence read files are uploaded via drag-and-drop, using on-the-fly compression by web-workers technology [18]. Compressed read files (.fastq.gz) are supported and recommended for fast upload. After upload, the user defines the condition that each read file represents. Samples sharing the same condition name are interpreted as replicates. Samples representing the control condition are further specified by marking a corresponding checkbox. The definition of at least one control sample is required. PinAPL-Py does not restrain the number of conditions that can be analyzed in one run. Next, the user selects the sgRNA library used in the screen from a drop-down menu. Currently, PinAPL-Py supports a set of 19 pre-set libraries, including the most common mouse and human genome-wide knock-out libraries such as GeCKO [19], or Brie/Brunello [17]. In addition, custom libraries can be uploaded. Finally, the user can adjust analysis options, such as methods for read count normalization, gene ranking, or various technical parameters for individual steps of the workflow. A manual can be found on PinAPL-Py’s documentation page.

### Comparison to other tools

We compared PinAPL-Py to MAGeCK and CRISPRcloud. A dataset from an unpublished CRISPR/Cas9 toxin resistance screen was used. Knock-out of the heparan-binding epidermal growth factor (*HBEGF*) is expected to confer resistance to the toxin used, making this gene a positive control for the comparison. A comparison to caRpools was not possible since the tool could not be run successfully and support was no longer available. Unlike both MAGeCK and CRISPRcloud, PinAPL-Py does not require the user to know the sequence composition of the reads as it extracts the 20 bp sgRNA sequence through automatic adapter alignment without input from the user. In addition to higher user convenience, adapter alignment also makes read identification robust against indels lying 5’ of the sgRNA (Fig. 3A, green arrow) and enables staggered read layout (Fig. 4). Another advantage of PinAPL-Py’s alignment protocol is the provision of both a matching threshold parameter and an ambiguity threshold parameters which allow the user to adjust the stringency of the read identification. Increasing stringency allows to safely discard reads matching multiple library sequences, thus avoiding the count of ambiguous, poorly designed sgRNAs (Fig 2; Fig 3A, red arrow) which can lead to false positive signals in the screen. On the other hand, decreasing stringency can be a useful means to control for non-optimal sequence quality.

**Fig 4:**
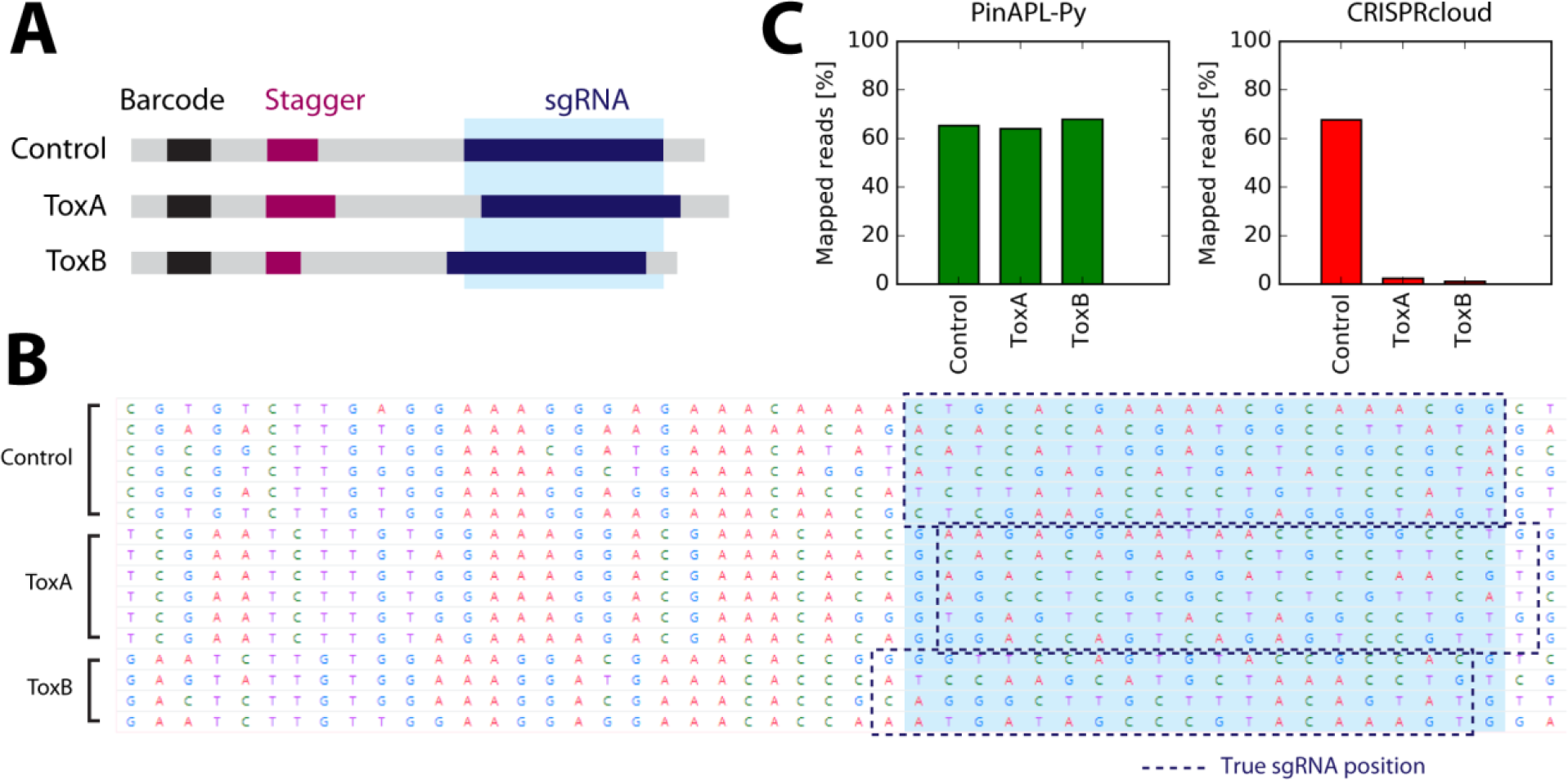
Alignment of staggered reads. **(A):** Staggering introduces spacer sequences of variable length (red) to cause shifts in the sequenced region, thereby enhancing nucleotide diversity per position when samples with different stagger lengths are pooled on the same sequencing lane. **(B):** Screenshot of CRISPRcloud’s read extraction page showing the framed area in (A). The definition of the 20 bp sgRNA sequence (light blue highlight) needs to be defined for all samples at once but misses the sgRNA region by +1 or −1 bp in the ToxA and the ToxB sample, respectively, due to the staggered read layout. **(C):** Since CRISPRcloud’s read identification does not to allow mismatches, read mapping fails in the corresponding samples.

#### Comparison to MAGeCK

As laid out above, PinAPL-Py requires less know-how from the user and offers more flexibility for read identification. Apart from differences caused by the processing of ambiguous reads, read counts, obtained with either PinAPL-Py or MAGeCK, are nearly identical (Fig. 3A, left panel; Fig 3B, left and middle panel). sgRNA fold-changes, as reported in either PinAPL-Py’s or MAGeCK’s enrichment analysis, show a high correlation and a large overlap of significant sgRNAs (Fig. 3C, left panel). *HBEGF* was retrieved with high significance (FDR < 0.005) as the top ranking gene by both PinAPL-Py and MAGeCK (Fig. 3D, left panel), and the ranked gene lists show good overlap (Fig. 3E). Finally, as mentioned above, MAGeCK has served as a highly appreciated analysis workflow in the field, but its usage is constrained by the user’s skill and computer platform. PinAPL-Py’s addresses this challenge by providing a user-friendly web-interface that can be run without any bioinformatic expertise. We conclude that PinAPL-Py constitutes an improvement over MAGeCK as its alignment protocol is more rigorous and flexible, and its web-interface makes sequencing analysis more easily accessible, while its functionality maintains the previously established standards.

#### Comparison to CRISPRcloud

CRISPRcloud also runs as a web-service, similar to PinAPL-Py. As with MAGeCK, read counts and sgRNA fold-changes, obtained with PinAPL-Py or CRISPRcloud, are almost identical, apart from differences caused by PinAPL-Py’s filtering procedure (Fig. 3A-C, right panels). CRISPRcloud and PinAPL-py differ in the way sgRNA results are combined into a ranking of genes. In particular, CRISPRcloud introduces novel gene scores (“directionality”, “hit ratio” and “conflict”) providing useful information about efficacy and consistency of the set of sgRNAs targeting each gene. However, these scores are fractional notions with number of sgRNAs per gene (which is typically no larger than 10) in the denominator, so they do not provide sufficient resolution to effectively sort a totality of 20,000 genes. As a consequence, *HBEGF* could not be retrieved as an outstanding positive hit by means of these scores alone. Taking the average log fold-change score into account as an additional criterion (see “Benchmarking & Quality Control”) yielded comparable results to PinAPL-Py’s gene ranking score (Fig. 3D,E). However, a gene ranking score like αRRA is more robust to single sgRNA outliers which is why CRISPRcloud’s top 25 gene list is slightly biased towards genes with fewer sgRNAs gaining significant enrichment (Fig. 3E, red fractions), and with fewer sgRNAs overall (Fig. 3E, arrow). As a significant advantage, PinAPL-Py allows processing of data involving staggered reads. Since PinAPL-Py’s adapter removal is based on sequence alignment, the position of the 20 bp sgRNA sequence can be flexible, and staggered reads can be correctly processed. In contrast, staggering is incompatible with CRISPRcloud as its workflow requires all reads to have the 20 bp sgRNA sequence at a fixed position across all samples (Fig. 4). Finally, PinAPL-Py makes no restrictions to the number of experimental conditions to be analysed in the same run, whereas CRISPRcloud is limited to five groups at a time. We conclude that PinAPL-Py constitutes an improvement over CRISPRcloud because of its higher flexibility and increased functionality in read alignment and gene ranking while providing an equally user-friendly web-interface.

## Conclusions

PinAPL-Py provides an analysis workflow for CRISPR/Cas9 screens that combines comprehensive functionality and established methods with an easy-to-use web-interface. It leverages state-of-the-art methods for the analysis of reads, quality control of the data and statistical scoring of sgRNA and genes. PinAPL-Py was designed to be used by the bench biologists who conduct the screens and receive raw sequencing results from their facility. It provides an easy, scalable, and versatile application to quickly control the quality of the screen and identify hits for replication and functional validation.

## Declarations

### Ethics approval and consent to participate

Not applicable

### Consent for publication

Not applicable

### Availability of data and materials

The datasets used and/or analysed during the current study are available from the corresponding author on reasonable request.

### Authors’ contributions

Workflow development: PS. Web-service implementation: TB/JK. Experimental data: RW/PS. Supervision (Experimental data): JE. Supervision (Workflow development): NL/OH. Manuscript: PS/NL/OH. All authors read and approved the final manuscript.

## Acknowledgements

We would like to acknowledge the valuable feedback from the participants of the CRISPR screening workshop organized by the San Diego Center for Systems Biology in October 2016 as well as the members of the UC San Diego CRISPR screening discussion group. In particular, we would like to thank Kristen Jepsen and her team at the UCSD Institute for Genomic Medicine.

